# Topical administration of novel FKBP12 ligand MP-004 improves retinal function and structure in retinitis pigmentosa models

**DOI:** 10.1101/2024.10.17.618826

**Authors:** Araceli Lara-López, Klaudia Gonzalez-Imaz, María Rodríguez-Hidalgo, Miren Sarasola- Gastesi, Leire Escudero-Arrarás, Santiago Milla-Navarro, Pedro de la Villa, Maialen Sagartzazu-Aizpurua, José Ignacio Miranda, Jesús María Aizpurua, Adolfo López de Munain, Ainara Vallejo-Illarramendi, Javier Ruiz-Ederra

## Abstract

**Purpose:** This study evaluates the therapeutic potential of MP-004, a novel FKBP12 ligand, in the treatment of inherited retinal dystrophies (IRDs). MP-004 targets FKBP12/RyR interaction, which is disrupted in several neurological disorders with underlying oxidative stress.

**Methods:** The toxicity and efficacy of MP-004 were examined *in vitro* in 661W cells. Efficacy was evaluated in phototoxic and H_2_O_2_-induced damage using impedance assays, calcium igaing and *in situ* PLA. *In vivo,* MP-004 efficacy was evaluated in the *rd10* mouse model of retinitis pigmenetosa (RP) by topical ocular instillation. Retinal function was assessed by electroretinography (ERG), visual acuity was measured using a water maze test, and retinal structure was analyzed morphometrically.

**Results:** MP-004 exhibited low toxicity (LD_50_: 1.22 mM) and effectively protected 661W cells from phototoxicity (EC_50_: 30.6 nM). Under oxidative stress conditions, MP-004 preserved FKBP12.6/RyR2 interaction, partially restored endoplasmic reticulum calcium stores and prevented cell death. *In vivo,* MP-004 significantly preserved retinal function in *rd10* mice, with ERG wave amplitude increases of up to 50% in scotopic and 71% in photopic conditions, corresponding to rod and cone functions, respectively. Additionally, MP-004 improved visual acuity for low spatial frequency patterns, and preserved retinal structure with a 23% increase in outer nuclear layer thickness, and preservation in the number of rods and cones and their segment length.

**Conclusions:** MP-004 shows promise as a therapeutic agent for RP, preserving retinal structure and function, likely through modulation of FKBP12.6/RyR2 interaction. Further studies are needed to explore its pharmacokinetics and efficacy in other IRD models.

## Introduction

Retinitis pigmentosa (RP) is the most common hereditary retinal dystrophy (IRD) and a leading cause of non-preventable blindness worldwide. Affecting approximately 1 in 4,000 people, RP is linked to mutations in over 80 genes, leading to the progressive loss of retinal photoreceptors and irreversible vision loss^1,2^. Despite significant advances in identifying new genes and understanding the molecular mechanisms underlying RP, many aspects of photoreceptor degeneration remain unclear^3,4^. Moreover, like other IRDs, RP currently lacks an effective standardized treatment for most of the patients^1,5^. Indeed, while gene therapy has shown promise in early stage IRDs, its application remains limited by the need to target specific mutations or genes^6^. So far, Luxturna (Novartis Europharm Limited) is the only product approved for the treatment of Leber congenital amaurosis caused by mutations in RPE65 gene, which affects around 1 in 81.000 people^7^.

Photoreceptor death in IRDs, including RP, is driven by multiple pathways, such as the activation of protein kinase G (PKG), calpains, the parthanatos pathway, and apoptosis mediated by caspases and endoplasmic reticulum (ER) stress^8–11^. Many mutations in IRD affect genes related to the phototransduction cascade, frequently causing dysregulation of cGMP, which has been known to be toxic to photoreceptors^12^. High cGMP levels activate PKG and CNG channels, increasing intracellular calcium and triggering calcium-activated calpains, especially calpain-1 and calpain-2, with distinct roles in neuroprotection and neurodegeneration, respectively^12–15^.

Extensive calpain activation is observed in several RP models, including the *rd1*, *rd2*, and *rd10* mice, as well as other animal models of photoreceptor degeneration^16^. In the retina, dynamic intracellular calcium fluxes are crucial for visual signal transduction, particularly in photoreceptor outer segments, where cytosolic [Ca^2+^] is essential for adapting and modulating the phototransduction cascade^17^. Calcium homeostasis is primarily regulated by the cGMP-activated cation channel (CNGC) and the Na^+^⁄Ca^2+^-K^+^ exchanger (NCK)^18^. However, ER and mitochondria also regulate the intracellular concentration of calcium through its release and storage^19^. The main ER channels involved in calcium homeostasis are ryanodine receptors (RyRs) and inositol 1,4,5-trisphosphate receptors (IP3R), which release calcium from the ER lumen into the cytoplasm, and the endo/sarcoplasmic reticulum calcium transporter ATPase (SERCA), which transports calcium back into the ER^20,21^. In pathological conditions, uncontrolled calcium leakage from ER channels may activate cytotoxic pathways in the mitochondria, including reactive oxygen species production, autophagy, and apoptosis^22,23^. Three RyR isoforms (RyR1-3) are expressed in the retina^24^. RyR2 has been implicated in photoreceptor death in a mouse model of achromatopsia^25,26^ and identified as a key factor in calcium dysregulation and photoreceptor death in CNGC-deficient models. In these models, deletion of RyR2 was found to reduce ER stress and mitigated photoreceptor loss^25^. Additionally, RyR2 overexpression has been linked to worsened photoreceptor degeneration through ER stress and the cGMP pathway^21^.

RyR channels are regulated by a complex system involving various accessory proteins, including FK506 binding proteins FKBP12 and FKBP12.6, which act as molecular switches to ensure proper channel function by stabilizing the closed state of RyR channels, preventing subconductance states, and facilitating coupling between adjacent channels^27,28^. Congenital mutations or post-translational modifications caused by nitro-oxidative stress may result in dysfunctional leaky RyR channels^29,30^.

Oxidative stress, resulting from the overproduction of reactive oxygen species (ROS), is a significant contributor to photoreceptor cell death in RP, particularly during the secondary cone degeneration that follows rod loss. The role of oxidative stress in photoreceptor death is well-documented across various models of retinal degeneration^31–34^ with studies highlighting the protective effects of antioxidants and the role of NADPH oxidase, a key enzyme in ROS production activated by elevated cytosolic calcium levels^35^. Light-induced ROS production has been shown to accelerate retinal degeneration in both inherited and age-related retinopathies, including RP^31,33^.

To address the broader challenge of premature photoreceptor death in IRDs, we have tested the effect of MP-004, a novel FKBP12 ligand, in cellular and animal models of RP. MP-004 belongs to a family of triazole molecules that potentiate FKBP12/RyR interaction and normalize intracellular calcium levels under oxidative stress conditions^36,37^. Our strategy aims to modulate a key pathogenic mechanism that is disrupted in most, if not all, IRDs, offering potential benefits regardless of the underlying genetic cause.

## Methods

### Preparation of MP-004

MP-004, a highly water-soluble salt optimized for ophthalmic applications, was prepared from compound MP-002 (formerly known as AHK2) which was synthesized following a CuAAC “click” methodology from 2-azidoethyl-N,N-dimethylamine and 4-methoxyphenyl propargyl sulfide^37^. To a 0.2M solution of the resulting 1-[2-[2-[2-(N,N-dimethylamino)ethyl]-4-[(4-methoxyphenyl)thiomethyl]-1*H*-1,2,3-triazole (MP-002) in anhydrous MeOH, cooled to 0 °C, was added 1.0 equiv of chlorotrimethylsilane and the mixture was stirred for 30 min at the same temperature. Evaporation of the solvent under reduced pressure and crystallization from acetone afforded pure non hygroscopic MP-004 (87%).

### Impedance Assays

Impedance was used to evaluate cell cytolysis in efficacy and toxicity assays. 661W mouse photoreceptor cells^38^ were kindly provided by Prof. Muayyad Al-Ubaidi (University of Oklahoma). 50,000 cells/well were seeded onto ECM-coated CytoView-Z 96-well impedance plates (#Z96-IMP-96B, Axion Biosystems) and grown in DMEM (#11594486, Fisher Scientific) with 40µl/L hydrocortisone 21-hemisuccinate sodium salt (#H-2270, Sigma), 40µl/L progesterone (#P-8783, Sigma), 0.032g/L Putrescine dihydrochloride (#P-7505, Sigma), 40µl/L β-mercaptoethanol (#M-6250, Sigma)^39^ at 37°C, 5% CO_2_. Cell cytolysis was evaluated by impedance measurement (1 min intervals, 10 kHz) using the Maestro Edge system (Axion Biosystems).

### Calcium imaging

Calcium imaging was used to evaluate MP-004 activity in normalizing cytosolic calcium levels in 661W photoreceptor-like cells under oxidative stress. 661W cells were seeded onto CytoView-Z 96-well plates at 50,000 cells/well, and after 2 days oxidative stress was induced by adding 2 mM H_2_O_2_ (#349887, Merck) for1 hour. The effect of MP-004 was evaluated at 1µM concentration. For basal cytosolic calcium levels measurement, cells were loaded with 4 µM Cal Red™ R525/650 AM (#20590, Deltaclon) for 30 minutes at 37°C and recordings were performed in imaging buffer (125 mM NaCl, 1.2 mM MgSO_4_, 5 mM KCl, 25 mM HEPES, 6 mM glucose, 2 mM CaCl_2_, pH 7.4) at room temperature, using a Glomax Discover microplate reader (Promega). ER calcium content was calculated using the fluorescence amplitude generated by adding 1 µM ionomycin in the absence of extracellular calcium. Intracellular calcium concentration was estimated by the ratio of Cal Red AM fluorescence intensities at 525 and 690 nm emission wavelengths, with excitation at 475 nm. Data were normalized to mean values of control cells.

### Ethics Statement and Animal Handling

All animal procedures were conducted following the ARVO Statement for the Use of Animals in Ophthalmic and Vision Research and approved by the Animal Care and Use Committee of Donostia University Hospital and the Clinical Research Ethics Committee of the Basque Country, Spain (CEEA16/013, 13 January 2017). *C57BL6/J* mice (wild-type, WT) were used as controls, and a congenic inbred strain of B6.CXB1-Pde6brd10/J mice in a *C57BL6/J* background (Pde6brd10, *rd10*), as a model of retinitis pigmentosa. Animals were obtained from Jackson Laboratory and were housed under a 12-hour light/dark cycle at 22°C, with 45-55% humidity and free access to food and water at the Biogipuzkoa facility. Mice received daily ocular instillations of 1.5 µl eye drops containing either MP-004 or vehicle (PBS) under inhalation anesthesia. MP-004 was administered for 4 days from P12 to P15 (15 µg, n=36), for 11 days from P14 to P24 with either 15 µg (n=32) or 30 µg (n=14), and for 16 from P14 to P29 (15 µg, n=9).

### Quantitative Real-Time PCR (qPCR)

Total RNA was extracted using the miRNeasy Mini Kit plus DNaseI (#217084, Qiagen), following the manufacturer’s instructions. cDNA was synthesized from 500 ng of RNA, using SuperScript Vilo cDNA Synthesis kit (#11754050, Thermo Fisher). qPCR was performed with 10 ng of cDNA and analyzed as previously described^40^, using the CFX384 system (BioRad), with Power SYBR Green PCR Master Mix (#4367659, Thermo Fisher). Samples were run in technical triplicates and gene expression was normalized to reference gene *Tbp*. Primer sequences for mRNA validation are shown in Table 1.

**Table 1.**
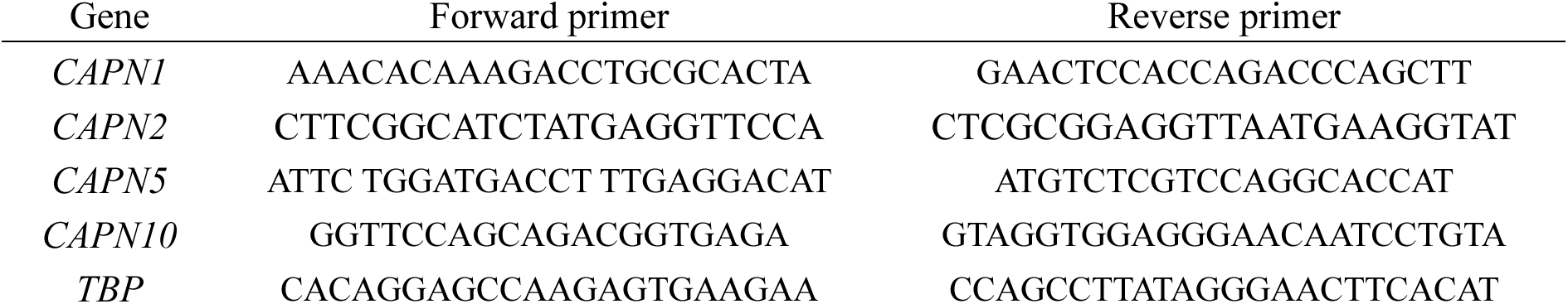
Primer sequences used for qPCR.

### Capillary Western Blot

Protein was extracted from WT and *rd10* mouse retinas using RIPA lysis buffer (50 mM Tris-HCl pH 7.2, 0.9% NaCl, 1% NP40, 1 mM EGTA, 1 mM EDTA) containing protease and phosphatase inhibitors (#748443, Thermo Fisher) and inhibitors of calpain I/II and cathepsins B/L (#208719, Sigma). Protein quantification was performed by Bradford protein assay (#500-0006, Bio-Rad). Capillary Western blot was performed using the Jess instrument (ProteinSimple, Biotechne), following the manufacturer’s instructions. Protein samples were diluted to 0.5-2 µg/µl with 5X Fluorescence Master Mix (Biotechne) and denatured at 95°C for 5 minutes. Samples were loaded onto customary plates (#SM-W004 or #SM-W005, Biotechne) for protein separation. Table 2 shows the list of antibodies used. Chemiluminescence signals were quantified using the Compass software, which generated chemiluminescence spectra and lane view images. Signals were normalized to total protein (#DM-TP01, Biotechne).

**Table 2.**
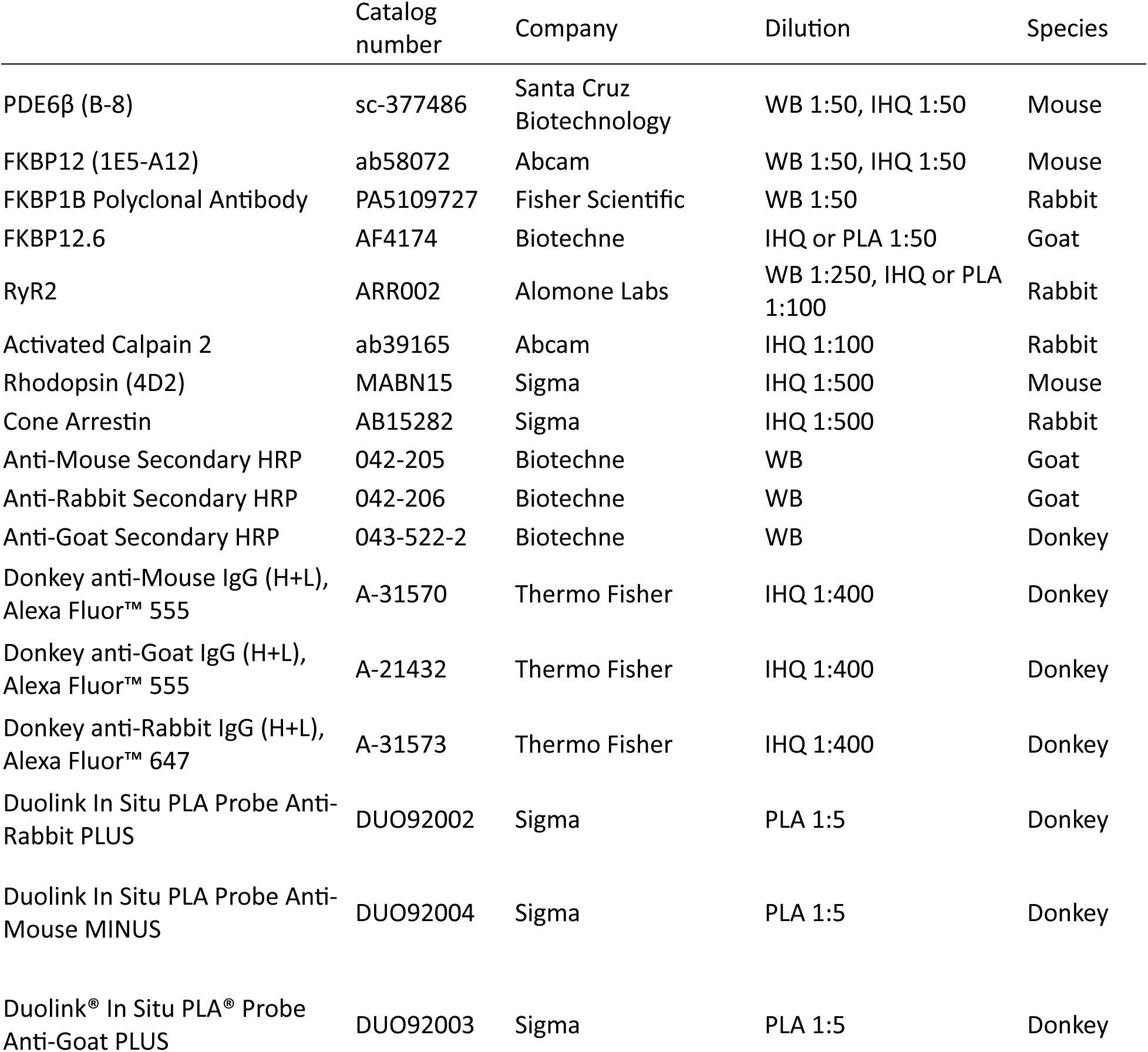
List of antibodies used in western blot (WB), immunohistochemistry (IHQ), and *in situ* proximity ligation assay (PLA) experiments.

### Immunohistochemistry

Immunohistochemical characterization was performed in retinal cryosections (7-12 µm) from P16, P25 and P30 mice, as previously described^41^. Briefly, sections were blocked for 1 hour in a solution containing 0.5% Triton X-100, 2% donkey serum, and 2% BSA in PBS and incubated with primary antibodies overnight at 4°C, followed by 1 hour-incubation of secondary antibodies and DAPI (#D9542, Sigma) at room temperature. Sections were mounted with Prolong Gold Antifade (#P36930, Thermo Fisher) and imaged by confocal microscopy (Zeiss LS900). Table 2 shows a list of antibodies used for immunohistochemistry (IHC).

### *In Situ* Proximity Ligation Assay (PLA)

PLA analysis was used for evaluating FKBP12/RyR2 and FKBP12.6/RyR2 interaction in 661W cells or retinal cryosections as previously described^36,37^. For cellular assays, 661W cells were seeded onto μ-Slide 15 Well 3D Glass Bottom (#81507, Ibidi) at 3000 cells/well. 1µM MP-004 treatment was added at seeding, and after 24 h, oxidative stress was induced by adding 2 mM H_2_O_2_ for 1 hour. Cells and retinal cryosections (7 µm) were incubated overnight at 4°C with RyR and FKBP12/12.6 antibodies (Table 2). The Duolink *in situ* PLA Far red assay kit (#DUO92013, Sigma) was used with corresponding conjugated probes (#DUO92002, #DUO92004, #DUO92003, Sigma, described in Table 2). Samples were mounted with ProLong Gold antifade reagent with DAPI (#P36962, Life Technologies, Eugene, OR, USA) and imaged with a Zeiss LS900 confocal microscope. Images were processed and analyzed using the ImageJ software (NIH) based on previous studies^42,43^. For retinal sections, local background in PLA images was removed in ImageJ by processing each image with a median filter followed by watershed technique to obtain individual dots, and particles between 0.1-10 µm^2^ were analyzed. For these samples, colocalization was analyzed in the outer nuclear layer (ONL) and photoreceptor segments (OS/IS) and for each localization data are represented as coverage (% of area of PLA signal/total area). At least 15 images per mice were analyzed. In cells, particles between 0.1-10 µm^2^ were directly analyzed and colocalization data are represented as particles/nuclei. At least 30 images per condition were analyzed with an average of 3 cells per image.

### TUNEL Assay

The terminal deoxynucleotidyl transferase-mediated dUTP nick-end labeling (TUNEL) assay was used to evaluate photoreceptor cell death in retinal sections, as previously described41. Retinal cryosections (7 µm) from P14 and P16 mice were evaluated using the DeadEnd Fluorometric TUNEL System (# G3250, Promega, Madison, WI, USA), following the manufacturer’s instructions. Slides were mounted with DAPI-containing medium for nuclear visualization and analyzed by confocal fluorescence microscopy (Zeiss LS900). The free ImageJ macro, TUNEL Cell Counter, was used for image analysis^44^.

### Electroretinogram Recordings (ERG)

Full-field electroretinography (ERG) was used to evaluate retinal function in P25 *rd10* mice and untreated littermates were used as controls. ERG protocols followed International Society for Clinical Electrophysiology of Vision guidelines and were performed as previously described^45^ with minor modifications. After overnight dark adaptation, anesthetized mice were placed on a 37°C heating pad, and mydriasis was induced with 1% tropicamide. Full-field flash ERG responses were recorded using a custom-made Ganzfeld stimulator, with increasing light flash intensities (-3 to 1 log cd·s/m²). Rod and cone mixed responses (b-mixed) were recorded under scotopic conditions. Responses were averaged, with intervals between flashes of 1.2 to 15 seconds. Cone responses (b-wave, photopic) were recorded after 5 minutes of light adaptation using 30 cd/m² background white light, with a 1.2 second interval between flashes. Scotopic and photopic responses were amplified using a NL104A pre-amplifier and NL100AK amplifier, (Digitimer LTD, Hertfordshire, UK.) and filtered using a band-pass filter set between 0.3 and 1000 Hz. All records were digitized using a PowerLab 4/30 (Adinstruments, Oxfordshire, UK).

### Visual Acuity Based on Water Maze

Visual acuity was assessed to evaluate visual function in P30 mice using a water maze, as previously described^46^. The test was conducted in a square tank (70 x 40 cm) filled with water (24–26°C), opaque on all sides except the front, where a screen was placed (Supplementary Figure S1). The screen was vertically divided into two equal parts: one half displayed a pattern of moving vertical bars, while the other half remained blank. The direction of the bars was randomized, with 50% moving to the right and 50% to the left, and the speed remained constant throughout the procedure. Mice were acclimated to the experimental conditions for one week prior to testing, learning to associate the area of the screen with the moving bars with the location of a submerged, hidden platform placed adjacent to that screen area. This platform, invisible under dim lighting, allowed the mice to escape from the water. During training, a high-contrast bar pattern (100%) with an optimal spatial frequency (0.088 cycles/degree) was used. In the experiment, the contrast and spatial frequency parameters were progressively adjusted until the mice could no longer detect the bar pattern^47^. Contrast sensitivity was determined by calculating the inverse of the minimum contrast at which the mice achieved a hit rate greater than 50%.

### Data Analysis

Data are expressed as mean or median ± standard deviation (SD), unless otherwise specified. Statistical analysis was performed using GraphPad Prism 8.3.0 and R-Studio^48,49^. Normality was assessed using the Shapiro-Wilk test and Q-Q plots. Normally distributed data were analyzed using t-tests or ANOVA followed by appropriate post hoc tests. For non-normal data, the Kruskal-Wallis test with Dunn’s post hoc test was applied. A hierarchical mixed-effects model was used for statistical analyses when a significant improvement in model fit was observed, as previously reported^50^. Pairwise comparisons were adjusted using the Bonferroni correction for multiple testing. In all cases, p-values < 0.05 were considered statistically significant.

## Results

### MP-004 protects from calcium dysregulation and 661W photoreceptor-like cell death induced by oxidative stress

MP-004 compound has previously shown activity in human myotubes under nitro-oxidative stress as a calcium normalizer by stabilizing FKBP12/RyR1 interaction^37^. Since oxidative stress is a major contributor to photoreceptor degeneration in RP^10^, we first wanted to evaluate the activity of MP-004 in rescuing photoreceptor cell death and calcium dysregulation induced by oxidative stress. To this end, we used H_2_O_2_ to induce oxidative stress in the 661W photoreceptor cell line. First, we assessed the median lethal dose (LD_50_) of H_2_O_2_ by performing life impedance measurements on 661W cells treated with 0-10 mM H_2_O_2_ for three hours. We found that the LD_50_ of H_2_O_2_ was 2 mM (Supplementary Figure S2), and, thus, this concentration was used to induce oxidative stress in 661W cells in subsequent experiments.

Next, we evaluated whether MP-004 was effective in reducing H_2_O_2_-induced toxicity in 661W cells. Indeed, Figure 1A shows a representative H_2_O_2_ cytolysis curve measured by impedance assays. Pretreatment of 661W cells under oxidative stress with 1 µM MP-004 resulted in reduced cytolysis over time compared to non-treated cells. In fact, MP-004 significantly reduced cell cytolysis from 50% in H_2_O_2_-treated cells, to 23.5% in H_2_O_2_ + MP-004, representing an overall cell death reduction of 53% (p<0.0001). Furthermore, while non-treated cells took 2.7 hours on average to reach 50% death, or kill time 50 (KT_50_), MP-004-treated cells took a significantly longer time with a KT_50_ of 3.8 h, representing a 35% increase in KT_50_ (p<0.0001, Figure 1B).

**Figure 1.**
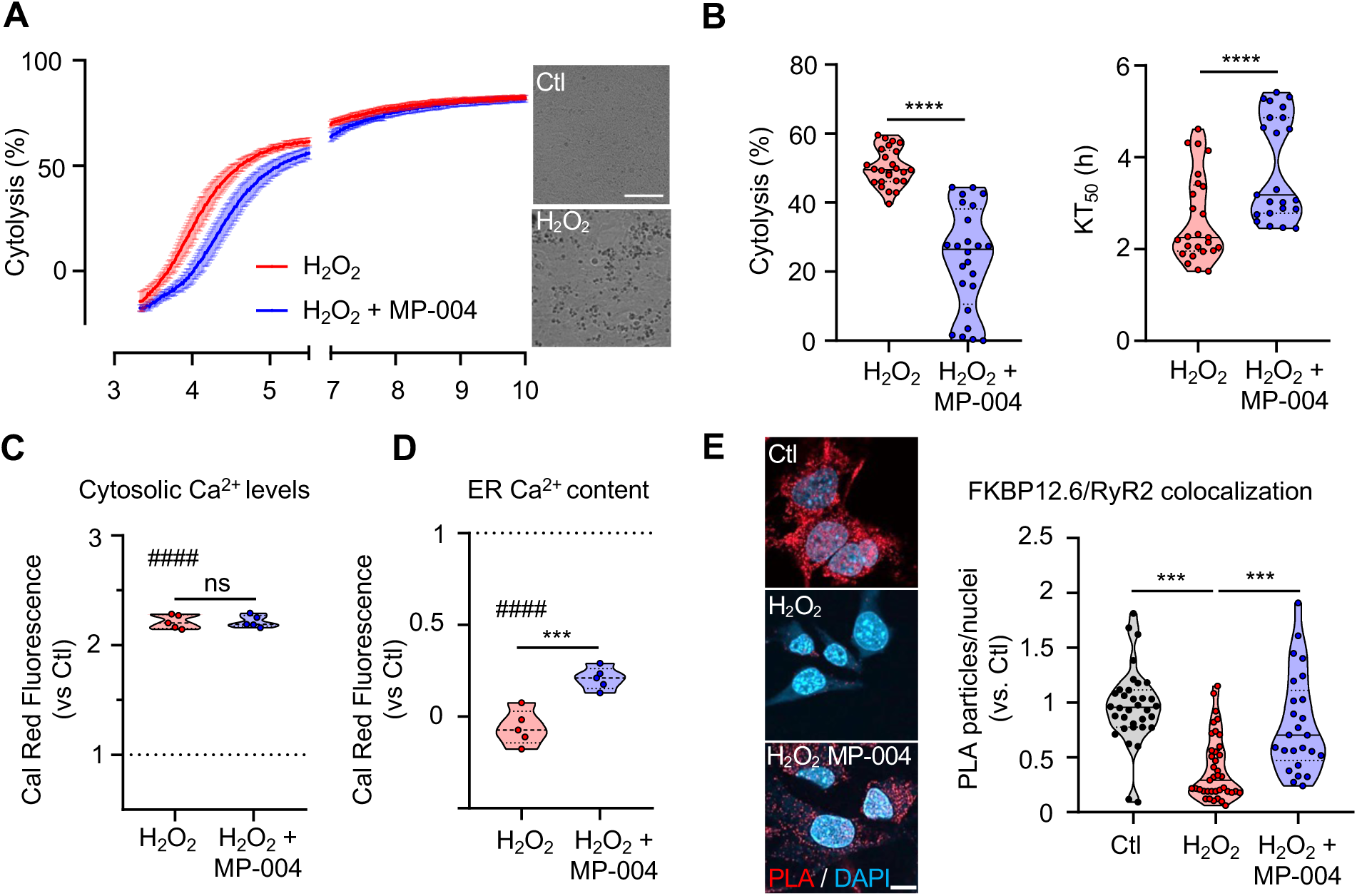
MP-004 prevents oxidative stress toxicity and increases ER calcium levels in 661W photoreceptor-like cells. **A)** Representative cytolysis time curve induced by 2 mM H_2_O_2_. Red shows non treated cells and blue shows cells treated with 1µM MP-004. Data expressed as mean ± SEM, n=5 wells/group. The right panel shows bright field images from cultures 6 hours after H_2_O_2_ addition. Scale bar: 100 µm. **B)** MP-004 protection against H_2_O_2_-induced cytolysis. KT_50_, kill-time 50. Dots represent individual wells from 5 independent experiments. ****p<0.0001, hierarchical mixed effects with pairwise comparisons and Bonferroni correction for multiple testing. **C)** Cytosolic calcium levels and **D)** ER calcium content measurements 1h after 2 mM H_2_O_2_ addition. Dots represent values from individual wells. ***p <0.001, one-tail unpaired t-test. ns, not significant. ^####^p<0.0001, one-tail unpaired t-test *vs.* control. **E)** FKBP12.6/RyR2 colocalization was quantified by in situ PLA. Left panel shows representative images of Control (Ctl) and cells under oxidative stress for 1 hour (2 mM H_2_O_2_). MP-004 effect was evaluated by overnight pretreatment (H_2_O_2_ + MP-004). PLA signal (red); nuclei (DAPI, blue). Scale bar: 10 µm. Right panel shows PLA quantification. Data are presented as violin plots with the median and interquartile range. Dots represent number of particles per nuclei of n=102 images from two independent experiments. ***p <0.001, hierarchical mixed-effects with pairwise comparisons and Bonferroni correction for multiple testing.

We next sought to evaluate whether MP-004 was able to normalize calcium levels in H_2_O_2_-treated 661W cells, by performing calcium life imaging with Cal Red fluorochrome (Figure 1C-D). Our data showed that 1 hour after addition of 2 mM H_2_O_2_ cytosolic calcium levels were significantly increased by 2-fold in 661W cells under oxidative stress compared to control cells (p<0.0001). However, pretreatment with MP-004 for 2 days did not rescue this cytosolic calcium elevation (Figure 1C). Next, we determined whether increased cytosolic calcium levels observed in photoreceptors under oxidative stress were associated with reduced calcium content in the ER (Figure 1D). Interestingly, we found that under oxidative stress, ER calcium content was drastically reduced in H_2_O_2_-treated cells (p<0.0001), which suggests that in 661W photoreceptor-like cells, oxidative stress induces calcium depletion from ER stores. Most importantly, pretreatment with MP-004 was able to partially rescue ER calcium content under oxidative stress, with a 25% recovery (H_2_O_2_ *vs.* H_2_O_2_ + MP-004; p = 0.0004) (Figure 1D).

Given that our results are consistent with leaky RyR channels as potential mediators of oxidative stress induced photoreceptor death, we next decided to evaluate whether FKBP12/RyR interaction was altered in 661W photoreceptor-like cells under oxidative stress. RyR2 is reported to be the major isoform expressed in retina^24^ and has previously been implicated in photoreceptor degeneration^25,26^. In the heart, it is generally accepted that RyR2 is stabilized mainly by FKBP12.6 protein, and that the loss of FKBP12.6/RyR2 interaction results in leaky RyR2 channels^29,51,52^. Therefore, we next wanted to evaluate FKBP12.6/RyR2 interaction in 661W cells under oxidative stress. To do this, we used the *in situ* PLA technique, which has been extensively used by our group and others to evaluate protein-protein interaction^53–55^, including FKBP12 with RyRs^36,37^. Our results show that upon H_2_O_2_ exposure, photoreceptors sustain a 40% loss of FKBP12.6/RyR2 interaction (p<0.0001, Figure 1E). Remarkably, this effect occurs 1 hour after addition of H_2_O_2_, well before the onset of cell death, and, therefore, it constitutes an early event within the oxidative stress cascade. Interestingly, pretreatment with MP-004 successfully prevented the loss of FKBP12.6/RyR2 interaction, as evidenced by a significant increase in their colocalization up to 80% (p<0.0001, Figure 1E). Overall, our results indicate that MP-004 protects 661W photoreceptor-like cells from oxidative stress death by stabilizing FKBP12.6/RyR2 interaction and normalizing ER calcium dysregulation.

### MP-004 presents a low toxicity profile and high efficacy in protecting 661W photoreceptor-like cells from phototoxicity

*In vitro* toxicity and efficacy of MP-004 was evaluated in 661W photoreceptor-like cells using impedance measurements. For toxicity dose-response curves, cells were exposed for 12 hours to seven doses of MP-004 ranging from 10 µM to 20 mM. MP-004 showed low toxicity, with an *in vitro* median lethal dose (LD_50_) of 1.22 mM (Figure 2A).

**Figure 2.**
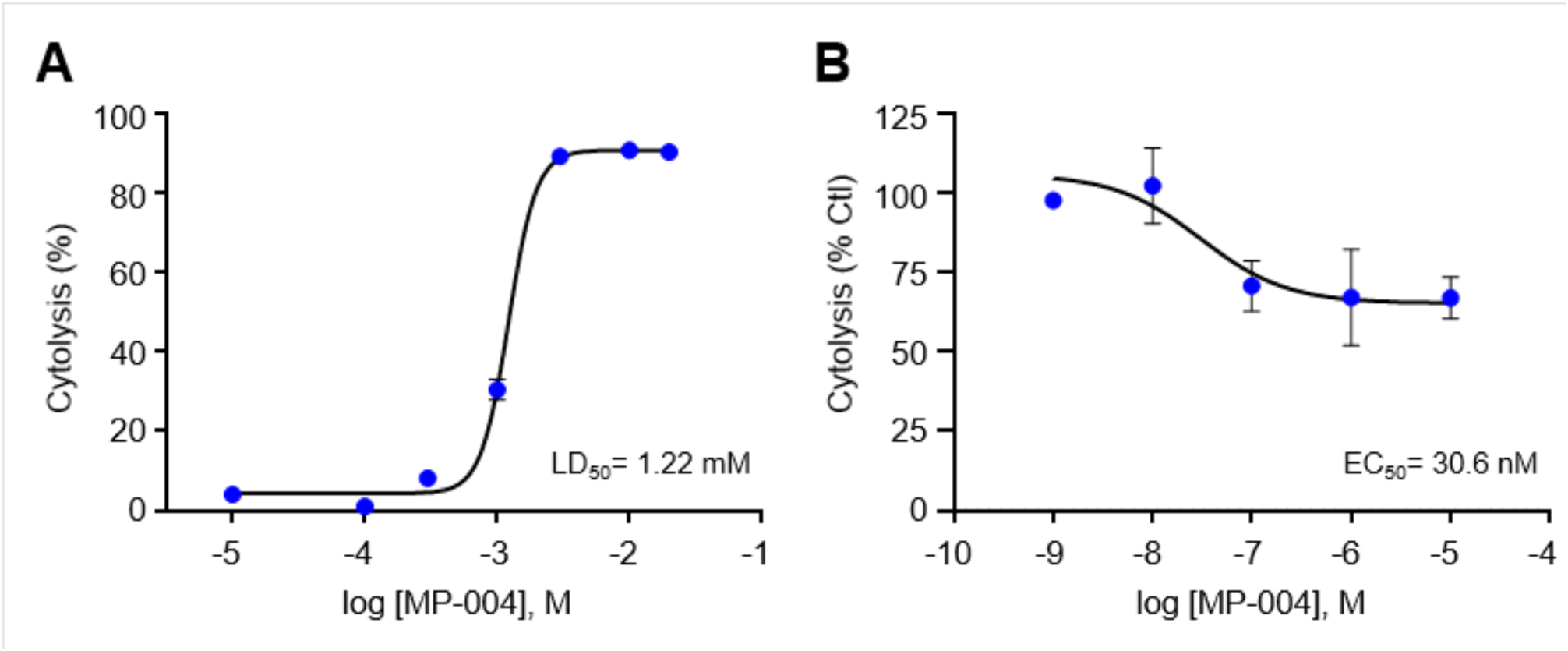
MP-004 *in vitro* toxicity and efficacy in 661W cells. **A)** MP-004 toxicity dose-response curve. LD_50_, MP-004 median lethal dose after 12 hours. Data are expressed as mean ± SEM, n=4 wells. **B)** Efficacy curve of MP-004 in a light-induced photoreceptor cell death model induced by 9-cis-retinal (20 µM) and white light (30,000 lux). MP-004 effect was measured after 6 hours and data are expressed as cytolysis percentage *vs*. non-treated. EC_50_, half maximal effective concentration of MP-004. Data are expressed as mean ± SEM from N=3 independent experiments performed in 3-6 replicates.

To analyze the *in vitro* efficacy of MP-004, we used a previously reported phototoxicity assay in which 9-cis-retinal and high-intensity white light were used to induce photoreceptor death due to conversion of 9-cis-retinal to all-trans-retinal^56,57^. In 661W cells, we found that after 6 hours of exposure, 20 µM 9-cis-retinal and 30,000 lux white light caused 45% cytolysis, while MP-004 protected 661W cells against 9-cis-retinal and light-induced phototoxicity in a dose-response manner with a half-maximal effective concentration (EC_50_) of 30.6 nM (Figure 2B). The maximum effect was achieved at 100 nM concentration of MP-004 and resulted in around 30% protection of phototoxicity. The *in vitro* therapeutic index of MP-004 derived from the toxicity (LD_50_) and efficacy (EC_50_) data was found to be 40,000, which indicates a wide therapeutic window and a minimal risk of toxicity at therapeutic doses.

### Disruption of FKBP12/RyR2 interaction in the retina of *rd10* mice

The expression and distribution of RyR2 and FKBP12 proteins have not been studied in the *rd10* mouse model, so we aimed to investigate these proteins in WT and *rd10* mouse retinas during the process of photoreceptor degeneration. Western blot analysis was performed to assess the expression levels of retinal proteins in WT and *rd10* mice at P12, P14, P16, and P25 (Supplementary Figure S3A). As expected, PDE6B, which is mutated in *rd10* mice, showed significant differences between WT and *rd10* samples, with a significant 60-80% reduction in the protein levels observed from P12 throughout the period analyzed. In contrast, no significant differences in protein levels were observed for RyR2, FKBP12 and FKBP12.6 between WT and *rd10* retinas. Likewise, immunohistochemistry studies (Figure 3A) did not reveal any differences in protein distribution for FKBP12, FKBP12.6, and RyR2 between WT and *rd10* at P16. These proteins are primarily localized in the outer and inner segments (OS/IS) of photoreceptors, with additional expression observed in the ONL (Figure 3A). To further investigate FKPB12/RyR2 protein interaction, we performed an *in situ* PLA in P14 and P16 WT and *rd10* mouse retinas. At P14, our analysis revealed a significant reduction in the ONL of FKBP12.6/RyR2 and FKBP12/RyR2 colocalization by 33% and 54%, respectively (Figure 3B). FKBP12.6/RyR2 colocalization showed mean values of 6.43 ± 1.78 (WT) and 4.32 ± 2.02 (*rd10,* p=0.03), and FKBP12/RyR2 showed mean values of 6.2 ± 2.45 (WT) and 2.87 ± 0.98 (*rd10,* p=0.013). Furthermore, FKBP12.6/RyR2 decreased interaction persisted at P16 in the photoreceptor segments (16.55 ± 2.94 in WT *vs.*11.5 ± 3.89 in *rd10*, p=0.04). Overall, our results indicate an early disruption of RyR2 interaction with FKBP12 and FKBP12.6 in the *rd10* mouse photoreceptors, prior to the onset of photoreceptor cell death.

**Figure 3.**
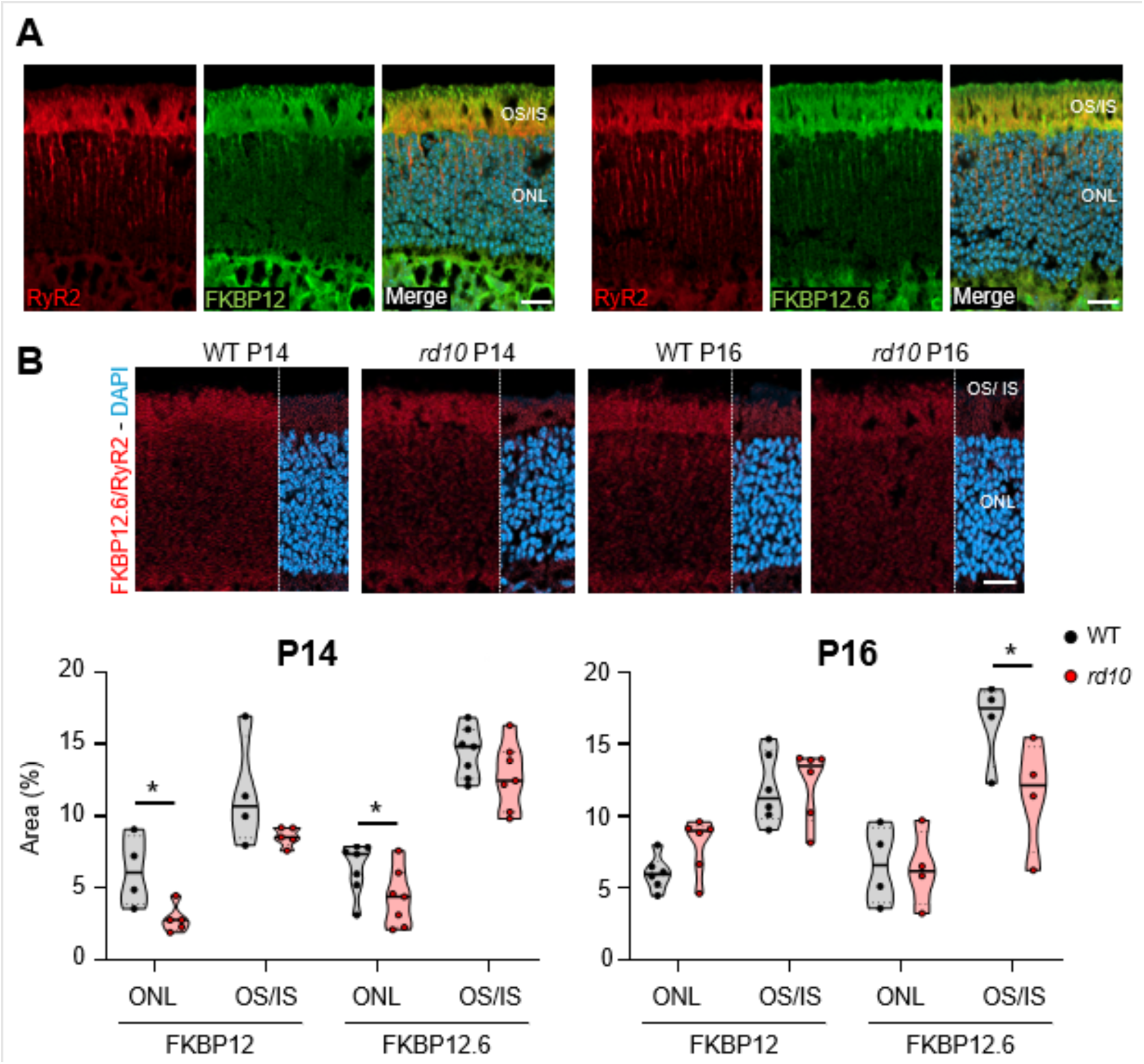
Characterization of RyR2, FKBP12, and FKBP12.6 in the mouse retina. **A)** Representative immunofluorescent images of P16 *rd10* retina sections. RyR2 (red), FKBP12 (green, left panel), FKBP12.6 (green, right panel), nuclei (blue). Scale bar 20 µm. **B)** In situ PLA analysis showing co-localization of FKBP12/RyR2 and FKBP12.6/RyR2 in retinas from WT and *rd10* mice at P14 and P16. Upper panels show representative immunofluorescent images of FKBP12.6/RyR2 colocalization. Red (PLA), nuclei (blue). Scale bar: 20 µm. The lower panel shows the quantification of PLA signals in the outer nuclear layer (ONL) and photoreceptor outer and inner segments (OS/IS). Dots represent mean values from individual mouse. At least 15 images were taken per retina. * p<0.05, one-tail unpaired t-test.

### MP-004 rescues photoreceptor cell death in *rd10* mice

To investigate the efficacy of MP-004 *in vivo*, we first assessed its biodistribution to determine if topical administration via ocular instillation would deliver therapeutically relevant concentrations to the retina. 18 µg of MP-004 were administered to WT mouse in 3 µL PBS eye drops and MP-004 concentration was measured in the retina by HPLC-MS over a 24-hour period (Supplementary Figure S4). The compound achieved retinal concentrations ranging from 0.5 to 1.5 µg/g, corresponding to an average concentration of 2.5 µM, which exceeds the effective concentration previously observed *in vitro*.

Once confirmed that MP-004 reaches the retina, we proceeded to study its effects on photoreceptor degeneration during the early stages of the disease. To this end, we treated *rd10* mice by ocular instillation with 15 µg/eye/day MP-004 from P12 to P15, and retinas were analyzed at P16. First, we verified that MP-004 had no significant impact on the protein levels of PDE6B, RyR2, FKBP12, and FKBP12.6 (Supplementary Figure S3B). Then, we examined the expression of calpains, a family of proteins largely implicated in photoreceptor degeneration in IRDs and other retinal disorders^12^.mRNA analysis revealed moderate increases of 20–30% in CAPN1, CAPN2, and CAPN5 expression in *rd10* retinas, with CAPN2 showing a significant 35% increase in *rd10* retinas compared to wild type (p < 0.05, Bonferroni). Treatment with MP-004 seemed to decrease CAPN2 overexpression although no statistical significance was achieved

Given that calpain 2 activity has been identified as a key event during rod photoreceptor cell death in the *rd10* model^13^, we conducted an immunohistochemical analysis to evaluate the influence of MP-004 on calpain 2 activation in *rd10* mice at P16. Similar to previous results at P18^13^, P16 *rd10* mice displayed a drastic elevation of active calpain 2 in the photoreceptor cells, with median numbers of activated calpain 2 positive cells in the ONL of 35 ± 20 cells/mm² for WT, 1577 ± 257 cells/mm² for *rd10* and 854 ± 1299 cells/mm² for treated *rd10* mice (Figure 4B). At P16, active calpain 2 distribution was restricted to the central retina, consistent with the characteristic central-to-peripheral degeneration gradient observed in this model^13,41^. Notably, neither the increase in calpain 2 activation in *rd10* retinas nor the 46% protection of MP-004 reached statistical significance, likely due to the high variability observed at P16 stage.

**Figure 4.**
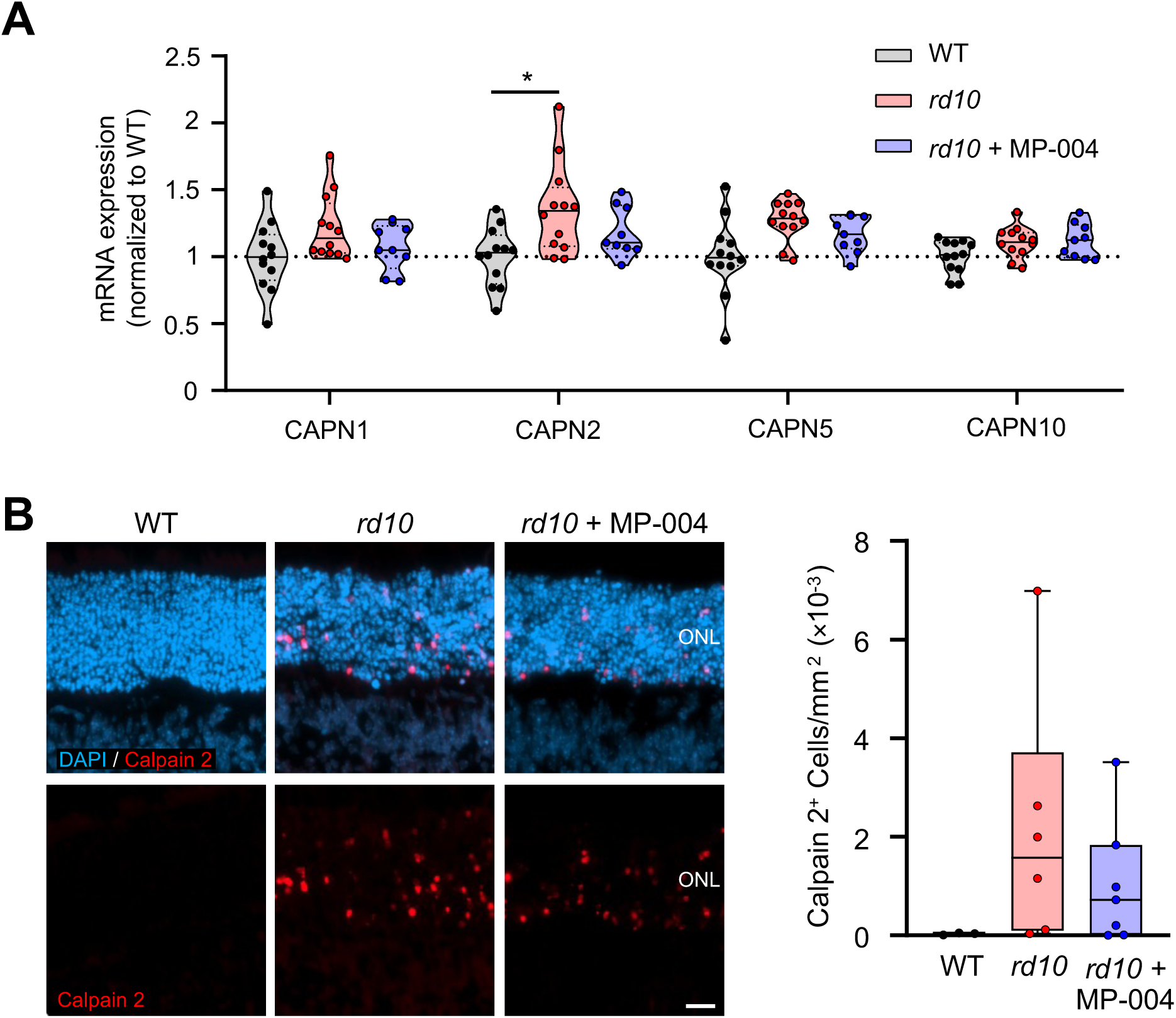
Effect of MP-004 on calpains in P16 *rd10* mice. **A)** mRNA expression levels of calpains in the mouse retina. *rd10* mice were treated daily with vehicle or MP-004 (15 µg/eye) from P12 to P15 via topical administration. Data are expressed as mean fold change over WT levels. Dots represent individual retinas. *p<0.05, hierarchical mixed effects with pairwise comparisons and Bonferroni correction for multiple testing. **B)** Immunohistochemical analysis of active calpain 2 in the *rd10* mouse retina at P16. Left panel shows representative immunofluorescent images of WT and *rd10* retina sections. Right panel shows quantification of the number of calpain 2 positive cells per mm^2^. Data are presented as violin plots showing the median and interquartile range. Dots represent values from individual mice. Kruskal Wallis analysis showed no statistical significance among the three groups. ONL, outer nuclear layer. Scale bar: 20 µm.

We then assessed photoreceptor cell death using the TUNEL assay at postnatal days P14 and P16, prior to the onset of degeneration and at an early stage of degeneration, respectively. At P14, both WT and *rd10* retinas exhibited low levels of TUNEL-positive cells in the photoreceptor layer, with means of 24 ± 14 and 34 ± 27 cells/mm², respectively. By P16, however, *rd10* mice showed a marked increase in TUNEL-positive cells, averaging 9,742 ± 5,410 cells/mm², compared to 206 ± 197 cells/mm² in WT mice, indicating significant photoreceptor loss (p<0.0001, Figure 5). At P16, TUNEL and calpain 2 positive cells showed a similar distribution with a predominant localization in the central retina. However, some TUNEL positive cells were also observed in mid-peripheral regions. MP-004 treatment provided a 56% protection against photoreceptor cell death in *rd10* retinas by reducing the number of TUNEL-positive cells to 4,421 ± 7,127 cells/mm² (p= 0.0006), which demonstrates its neuroprotective effect in the retina.

**Figure 5.**
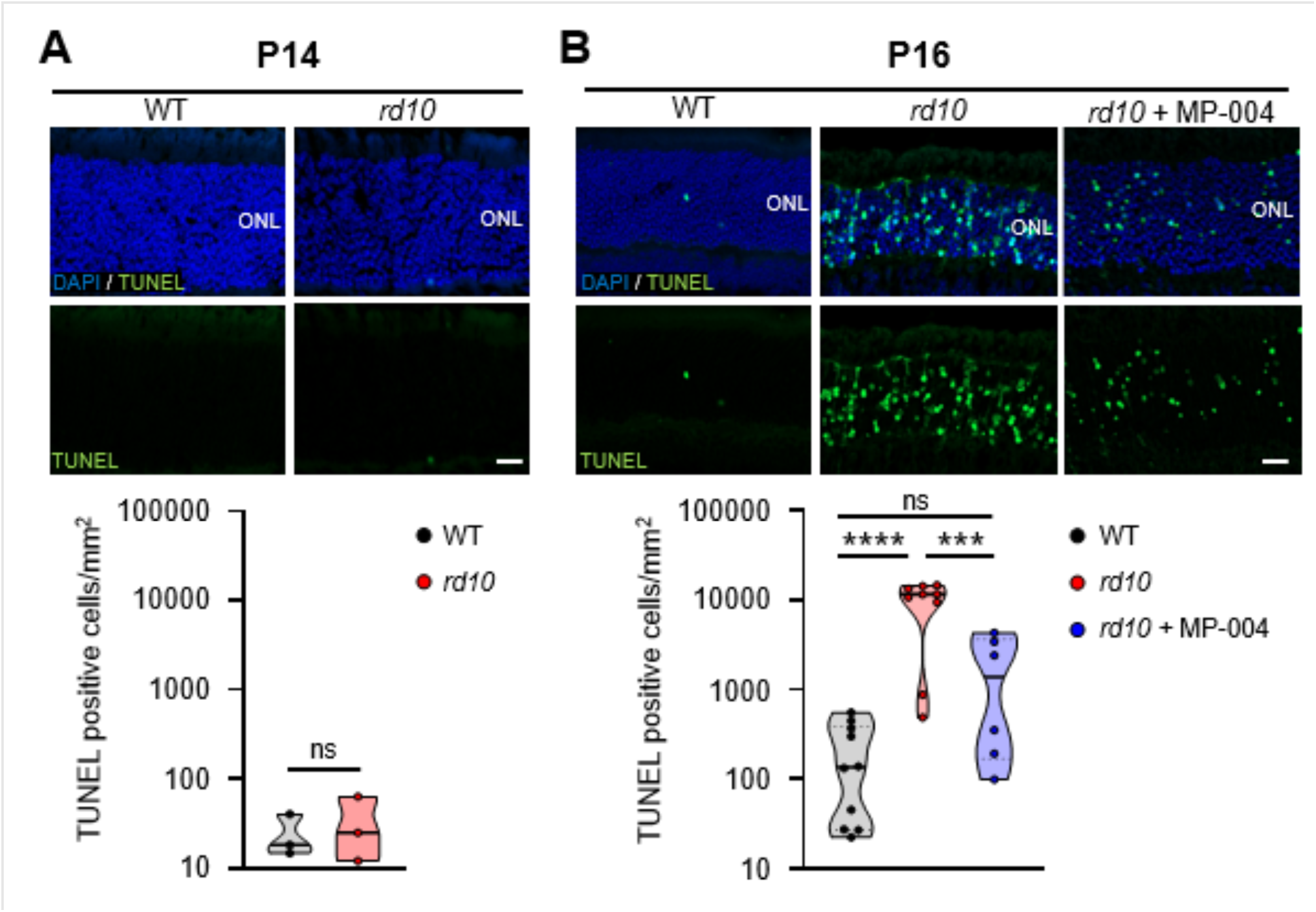
MP-004 reduces photoreceptor death in P16 *rd10* mouse. Analysis of cell death in P14 (A) and P16 (B) WT and *rd10* mice. Upper panels show representative images of TUNEL staining (green) and DAPI (blue). Scale bars: 20 µm. Lower panels show cell death quantification expressed as TUNEL positive cells per mm^2^. **A)** No differences were found at P14 between WT and rd10 mice. One-tail unpaired t-test. NS: non-significant. **B)** P16 *rd10* mice were treated with vehicle or MP-004 (15 µg/eye) for 4 days. Data are presented as violin plots showing the median and interquartile range. Dots represent values from individual mice. ***p<0.001, ****p<0.0001, One-Way ANOVA with Tukey’s post hoc test.

### MP-004 Enhances ERG Response, Visual Acuity, and Retinal Structure in *rd10* Mice

This study aimed to evaluate the effects of MP-004 on visual function in *rd10* mice during postnatal days 25 to 30 (P25-P30), a stage characterized by advanced photoreceptor degeneration^58–61^. Starting at P14, the mice received daily ocular instillations of MP-004 at 15 µg per eye.

ERG was conducted at P25 to assess rod and cone function under scotopic and photopic conditions, respectively. The b-wave response was used as an indirect measurement of photoreceptor activity. The mixed response representing the combined activity of both rod and cone photoreceptors (downward deflection), followed by a b-wave (upward deflection), indicating bipolar cell activity postsynaptic to the photoreceptors^62^.

As previously reported^63^, *rd10* mice exhibited a marked reduction in the amplitudes of mixed a/b-waves under scotopic conditions. However, MP-004 treatment led to a significant increase in b-wave amplitudes under mixed, and photopic conditions compared to vehicle-treated *rd10* mice (Figures 6 and 7). Specifically, ERG analysis revealed significant improvements in b-wave amplitudes in the retinas of MP-004-treated *rd10* mice. Under mixed conditions, b-wave amplitude increased by 50% (p=0.0125), 44% (p=0.0003) and 30% (p=0.0002) at stimulus intensities of 0.35, 0.67 and 1 log cd×s/m^2^, respectively with daily MP-004 treatment of 15 µg per eye (Figure 6B). Under photopic conditions, a 71% increase in b-wave amplitude was observed at -0.5 log cd·s/m² (p=0.0007), suggesting that MP-004 improves both rod and cone function (Figure 6C).

**Figure 6.**
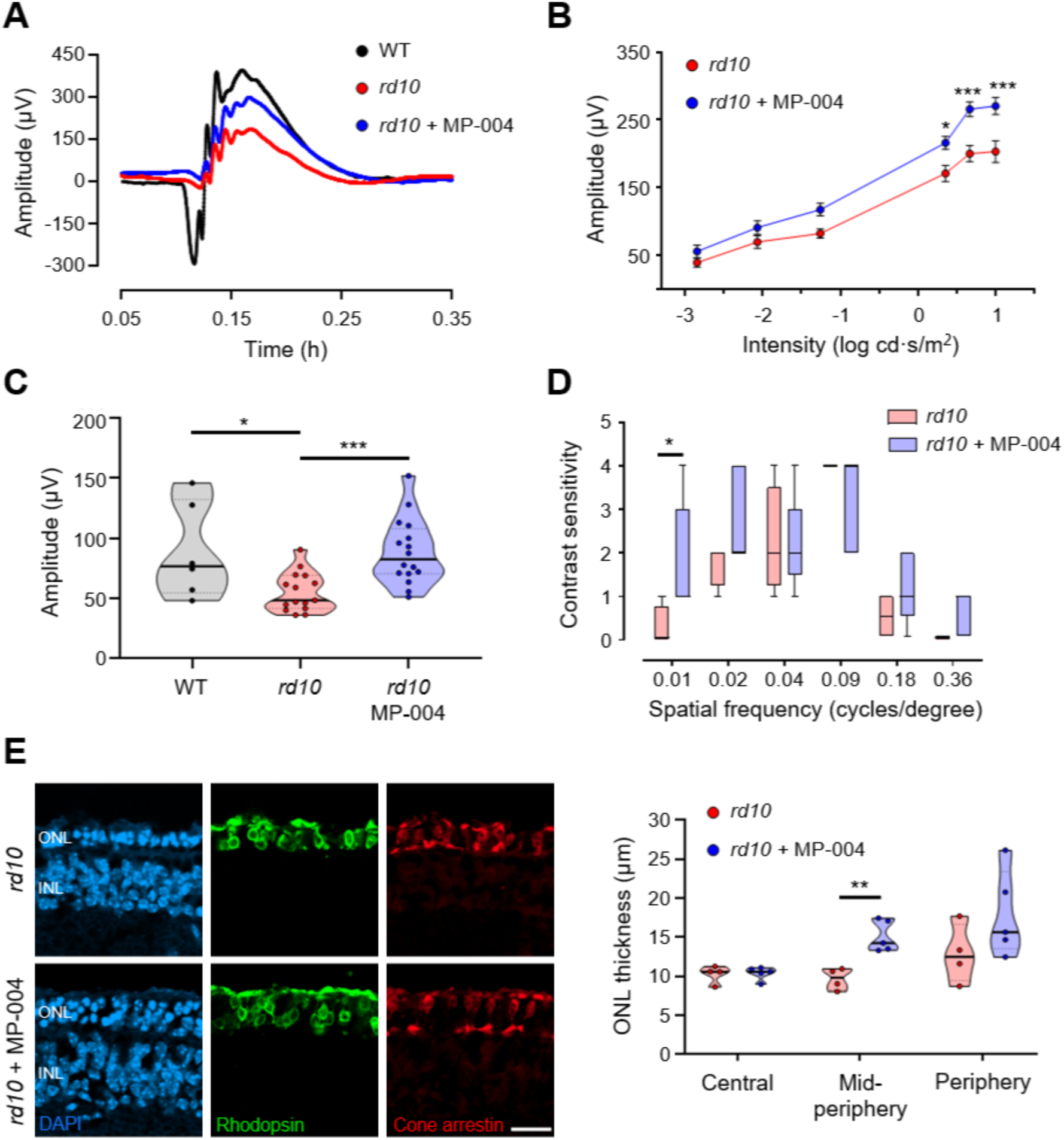
*In vivo* efficacy of MP-004 in *rd10* mice at a daily dose of 15 µg per eye. **A)** Representative ERG recordings from P25 WT (black) and *rd10* mice treated with MP-004 (blue) or vehicle (PBS, red) from P14 to P24. **B)** Intensity-response curves of mixed b-waves (scotopic) to light stimuli (-3 to 1 log cd×s/m^2^) in *rd10* mice treated with MP-004 or vehicle. Data are expressed as mean ± SEM. n=38 mice, *p<0.05, ***p<0.001, two-way ANOVA followed by Sidak’s post hoc test. **C)** Amplitude of cone responses (b-wave, photopic) to light stimuli (-0.5 log cd×s/m^2^) in WT and *rd10* mice treated with MP-004 or vehicle. n=37 mice, * p<0.05, ***p<0.001, Kruskal-Wallis test. **D)** Efficacy of MP-004 on visual acuity in P30 *rd10* mice. Data are represented as box and whisker plots showing the median and interquartile range. n=9 mice. * p<0.05, Mann-Whitney. **E)** Left, representative images retinal sections from P30 *rd10* mice showing nuclei (DAPI, blue), rhodopsin (green) and cone arrestin (red) distribution. Right, quantification of outer nuclear layer thickness. Data are presented as violin plot showing the median and interquartile range. Dots represent values from individual mice. **p<0.01, one-tail unpaired t-test. INL: inner nuclear layer; ONL: outer nuclear layer. Scale bar: 20 μm.

**Figure 7.**
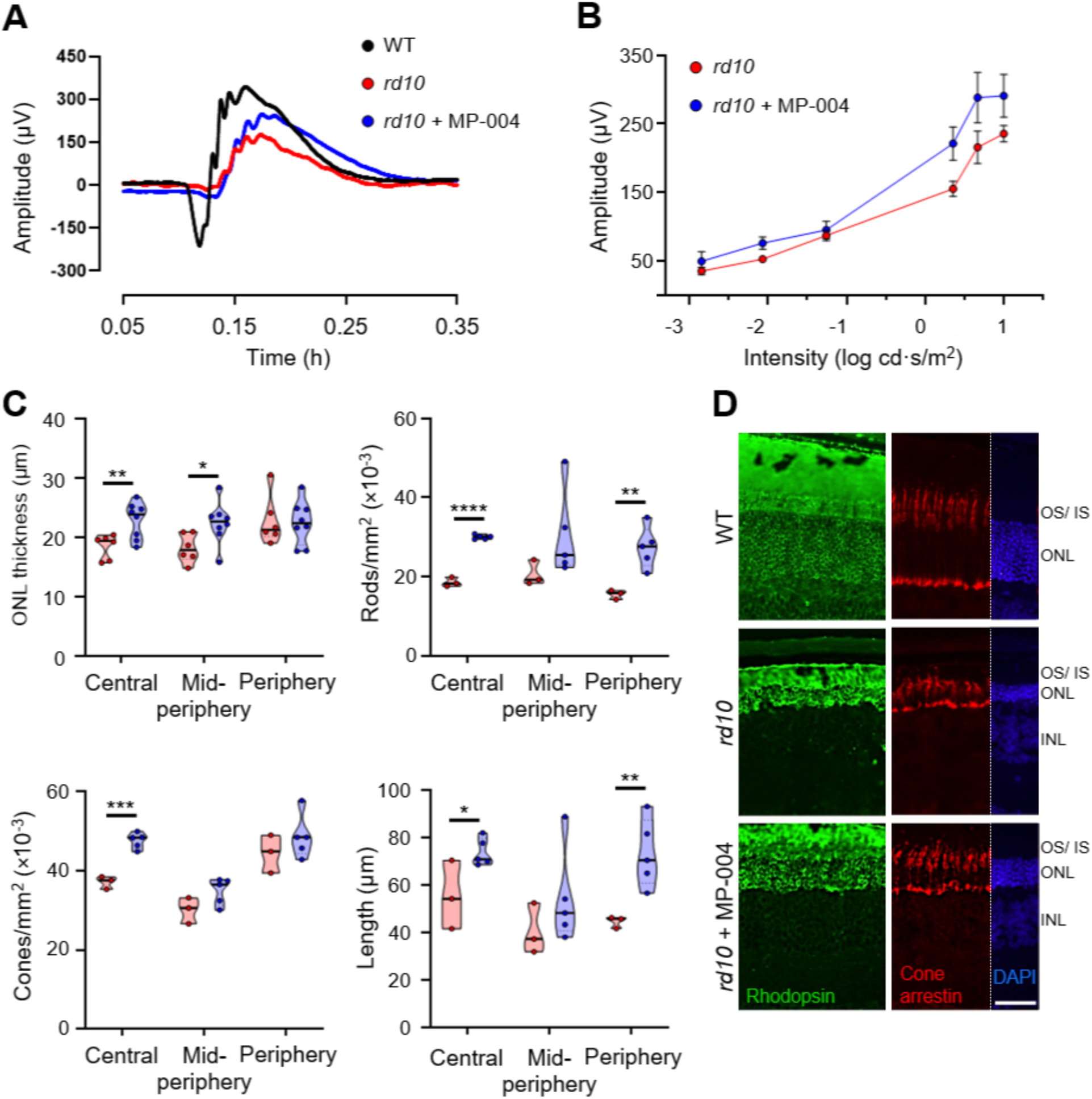
*In vivo* efficacy of MP-004 in *rd10* mice at a daily dose of 30 µg per eye. **A)** Representative ERG recordings from P25 WT (black) and in *rd10* mice treated with MP-004 (blue) or vehicle (red) from P14 to P24. **B)** Intensity-response curves of ERG b-waves to light stimuli (-3 to 1 log cd×s/m^2^) in *rd10* mice treated with MP-004 or vehicle. Data are expressed as mean ± SEM. n=14 mice. **C**) Morphometric analysis of P25 *rd10* retinas. Analysis of ONL thickness, number of rods and cones per mm^2^, and photoreceptor outer segment length. Data are represented as violin plots showing the median and interquartile range. Dots represent values from individual mice. *p<0.05, ** p<0.01, ***p<0.001, **** p<0.0001, one-tail unpaired t-test. **D)** Representative immunofluorescence images of P25 sections from WT, untreated and treated *rd10* animals showing nuclei (DAPI, blue), rhodopsin (green) and cone arrestin (red) distribution. INL: inner nuclear layer. ONL: outer nuclear layer. OS/IS: outer segment/inner segment. Scale bar: 20 μm.

To further determine whether the improvements in retinal function translated into enhanced visual acuity, a modified water maze test was performed at postnatal day 30 (P30), allowing for the required training period for the animals (Figure 6D). Contrast sensitivity was measured across a range of spatial frequencies (0.011–0.355 cycles/degree). A trend toward improved visual acuity was observed in MP-004-treated animals, with statistically significant differences at the lowest spatial frequency (0.011 cycles/degree, p<0.05), which represents one of the most challenging conditions. This suggests an improvement in low-contrast visual capacity.

Retinal morphometric analysis of P30 *rd10* mice revealed that MP-004 treatment led to a 56% increase in mid-peripheral retinal thickness (p= 0.0012, Figure 6E), indicating a protective effect on retinal structure. However, at postnatal day 25 (P25), no significant differences were observed between MP-004-treated *rd10* mice and untreated littermates in terms of the ONL thickness, or the number of photoreceptors (Supplementary Figure S5). Despite this, P25 *rd10* mice treated with MP-004 demonstrated significant improvements in rod and cone activity, as evidenced by our ERG results.

To further assess the efficacy of MP-004 on the preservation of retinal cytoarchitecture, we evaluated the effects of MP-004 at a higher dosage of 30 µg/eye/day. At the functional level, ERG analysis revealed similar effects of MP-004 to those observed in *rd10* mice treated with the lower dose of 15 µg/eye/day (Figure 7A, B). Specifically, ERG data demonstrated a significant increase in b-wave amplitude 33% (p=0.018), 42% (p=0.0006) and 37% (p<0.0001) at 0.35, 0.67 and 1 log cd×s/m^2^, respectively under mixed lighting conditions (Figure 7B).

More importantly, treatment of *rd10* mice with 30 /eye/day resulted in significant structural preservation of the retinal cytoarchitecture by P25. Morphometric analysis revealed a 24% increase in ONL thickness in the central retina (p=0.0103) and a 23% increase in the mid-peripheral retina (p= 0.0285) (Figure 7C, D). Furthermore, the number of rods was markedly preserved, with a 61% increase in rod cells in the central retina (p<0.0001) and a 76% increase in the peripheral retina (p= 0.0048). Cone cell numbers also showed an increase of 28% in the central retina (p= 0.0001). Moreover, MP-004 treatment at 30 µg/eye/day effectively preserved photoreceptor segment length, with a 33% increase in the central retina (p= 0.0198) and a 64% increase in the periphery (p= 0.0078).

These findings underscore the dose-dependent efficacy of MP-004, demonstrating substantial functional and structural benefits at higher doses.

## Discussion

This study presents a novel therapeutic approach for RP using MP-004, a new FKBP12 ligand known to stabilize FKBP12/RyR interaction and normalize calcium levels under nitro-oxidative stress^36,37^. In cellular models of oxidative stress, MP-004 significantly enhanced RyR2 and FKBP12.6 interaction, restoring ER calcium levels and preventing photoreceptor death under stress conditions. Additionally, in the *rd10* mouse model of RP, topical administration of MP-004 reduced photoreceptor death and preserved visual function and retinal architecture during the early stages of degeneration. These findings support MP-004 as a promising therapeutic candidate for RP and other IRDs.

In RP, oxidative stress is a major contributor to photoreceptor degeneration^10,31,32^. This oxidative stress can initiate a vicious cycle in which elevated ROS levels generate leaky RyR channels, which impact mitochondria, and further increases ROS production, ultimately leading to cell degeneration^64^. In 661W cells, we have shown that oxidative stress induces early calcium dysregulation, characterized by increased cytosolic calcium levels and depleted ER calcium stores. A previous study in 661W cells reported that sustained cGMP elevation leads to high intracellular calcium levels, ROS generation, and calpain activation^12,14,15,65^, suggesting a close interrelationship between calcium dysregulation and oxidative stress in IRDs.

Excessive calcium release from the ER is a well-documented trigger of photoreceptor death, largely due to mitochondrial dysfunction and activation of the parthanatos pathway and calpains^8,11^. In this study, we show that MP-004 significantly reduces cell death and replenishes ER calcium stores in oxidative-stressed 661W cells, suggesting that MP-004 has a protective role against H₂O₂-induced damage in photoreceptor cells. However, MP-004 failed to normalize elevated cytosolic calcium levels in 661W stressed cells. This could be attributed to the high concentration of H_₂_O_₂_ used (2 mM) and the fact that ROS affect not only RyR channels but also multiple calcium regulatory components such as Ca²⁺ transport at the plasma membrane, store-operated calcium entry, voltage-gated calcium channels, and SERCA and IP3 receptors in the ER^66^. Despite this, the partial restoration of depleted ER calcium stores by MP-004 indicates that RyR2 contributes to ER calcium depletion driven by oxidative stress.

In 661W cells, we found that oxidative stress disrupts the interaction between FKBP12.6 and RyR2, while MP-004 significantly preserves this interaction. These findings are consistent with our previous studies where MP compounds effectively restored FKBP12/RyR1 interaction and calcium homeostasis in human myotubes subjected to nitro-oxidative stress^36,37^. Interestingly, we found that the interaction between RyR2 and FKBP12/12.6 is compromised in the *rd10* mouse model at P14, well before major neurodegeneration occurs, hitting at an early role for RyR2 and impaired ER calcium signaling in RP. Other studies have linked RyR2 dysfunction to photoreceptor degeneration in IRDs^25,26^, reinforcing the hypothesis that disrupted ER calcium signaling is a common pathological mechanism in these diseases. These findings indicate that MP-004 could be beneficial in RyR2-mediated calcium dysregulation, in RP and possibly across a spectrum of IRDs.

Light-induced photoreceptor damage has been linked to mutations in genes associated with the visual cycle, phototransduction, and retinal health, such as *Rhodopsin*, *Abca4* , *Cralpb, Crb1, Eys*, *Mertk, peripherin/Rds, Rdh12*, as well as genes involved in melanin production^67–72^.

Phototoxicity significantly contributes to retinal degeneration through mechanisms that involve excessive ROS production and calcium dysregulation, ultimately leading to photoreceptor death and vision loss^73,74^. In RP, the interplay of oxidative stress and light exposure is particularly detrimental, accelerating the degeneration of cone cells following the loss of rod photoreceptors^10,75^. Several studies have shown that rearing *rd10* mice in darkness delays photoreceptor degeneration by up to 2 months^76^. Our findings in 661W cells demonstrate that MP-004 has a protective effect against light-induced phototoxicity with an effective concentration of 30.6 nM, highlighting its therapeutic potential for IRDs and other retinal diseases where light-induced damage is well-documented such as age-related macular degeneration and diabetic retinopathy^77,78^.

Our *in vitro* findings were confirmed in the *rd10* mouse model of RP, where MP-004 was administered daily via ocular instillation. A short 4-day treatment with MP-004 led to a 55% reduction in photoreceptor cell death, suggesting that MP-004 enhances photoreceptor survival by inhibiting calcium-mediated cell death. While calpain 2 activation is involved in calcium-induced cell death in *rd10* mice^8,79^, the high variability observed in P16 *rd10* mice limited definitive conclusions about its role in MP-004 photoreceptor protection. Notably, calpain 2 activation accounted for less than 20% of TUNEL-positive cells and its distribution was restricted to the central retina at P16, whereas TUNEL-positive cells were also present in the mid-peripheral retina, suggesting the involvement of other key players in early photoreceptor death in *rd10* mice. Future experiments will focus on studying the other cell death pathways, to further elucidate the MP-004 effect on photoreceptor protection.

The preservation of photoreceptors through topical MP-004 resulted in improved retinal function, as evidenced by ERG analysis that showed increased b-wave amplitudes, particularly in bipolar rod pathways, which are primarily affected in RP^63^. These functional improvements, along with enhanced visual acuity indicate that MP-004 activity extends beyond cellular protection, translating into meaningful physiological benefits. MP-004 treatment, particularly at higher dosage, preserved the retinal architecture which is crucial for sustaining visual function in RP in the long term.

MP-004 represents a promising advancement in the treatment of retinitis pigmentosa (RP) by selectively modulating the FKBP12/RyR2 interaction during oxidative stress. This targeted approach effectively addresses calcium dysregulation compared to conventional calcium channel blockers like diltiazem, which may lead to further calcium depletion and adversely affect photoreceptor survival^80^. Several therapeutic strategies for RP are currently being explored, including the administration of neurotrophic factors via an adeno-associated viral vector ^81^,

GSK-3 modulators^82^, and cGMP analogs that inhibit PKG^83,84^. While these strategies show potential, they are not without risks, as they may negatively impact phototransduction or carry additional risks associated with chronic systemic administration^83^. Ultimately, the future of RP treatment likely lies in combined therapies that simultaneously target multiple pathways to effectively combat this complex disease.

In conclusion, MP-004 represents a promising new approach for the treatment of RP, as it directly addresses calcium dysregulation and oxidative stress, two of the key drivers of photoreceptor degeneration in this disease. MP-004 demonstrates efficacy in preserving photoreceptor integrity in the *rd10* mouse model and in a photoreceptor-like cell line subjected to high oxidative stress or phototoxicity. This, in combination with its non-invasive topical administration, positions MP-004 as a promising therapeutic candidate not only for RP and IRDs, but also for more common ophthalmic diseases such as diabetic retinopathy or age-related macular degeneration. Further exploration of MP-004, in particular its biodistribution in larger animal models and its efficacy in other models of retinopathy, would significantly broaden its impact, potentially transforming the therapeutic landscape for both RP and other eye diseases with underlying oxidative stress and calcium dysregulation.

## Acknowledgements

The authors would like to thank Mª Carmen Sampedro from Servicio Central de Análisis Unidad de Álava, SGIker at UPV/EHU, and Laura Ramirez from the University of Alcalá for their excellent technical support. The authors also thank Carlo Manno from Rush University Medical Center and Maider Mateo-Abad from the Methodological Support Unit at IIS Biogipuzkoa for their expertise and assistance in statistical analysis, and Virginia Arechavala for kindly reviewing the final version of the manuscript. ChatGPT and DeepL Write were solely used for assisting in English correction and clarity of the manuscript.

---

**Supplementary Figure S1.**
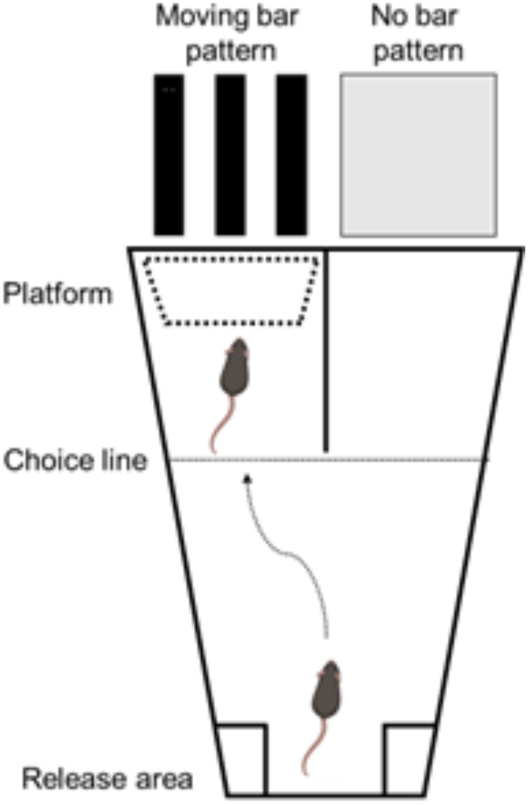
Schematic representation of the modified Morris water maze test used to evaluate visual acuity (adapted from Prusky et al., 2000).

**Supplementary Figure S2.**
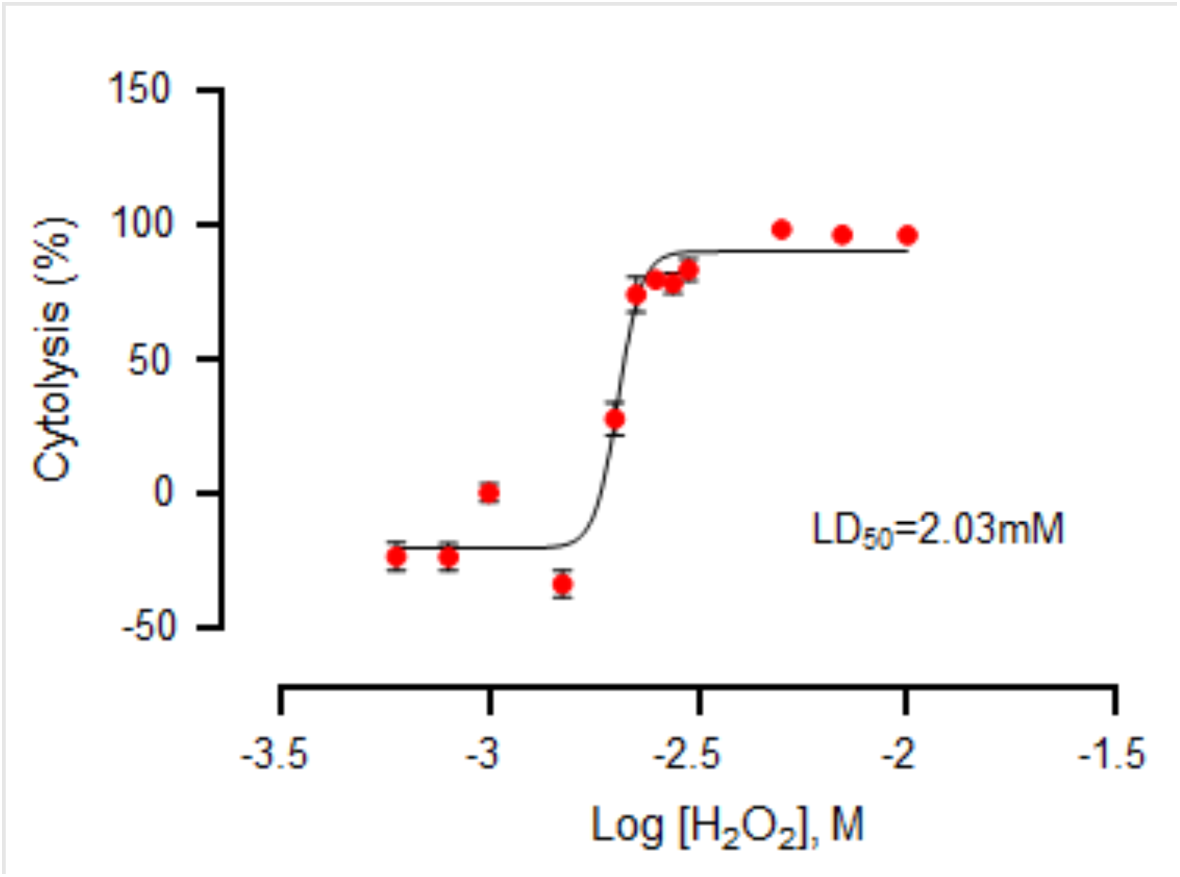
H_2_O_2_ toxicity dose-response curve in 661W cells. H_2_O_2_ toxicity was determined in 96-well plates using cell impedance assays. Data are expressed as mean ± SEM, each value is the average of n=5 wells. Data from one representative experiment out of three is shown. LD_50_, lethal dose 50%. erformed in 3-6 replicates.

**Supplementary Figure S3.**
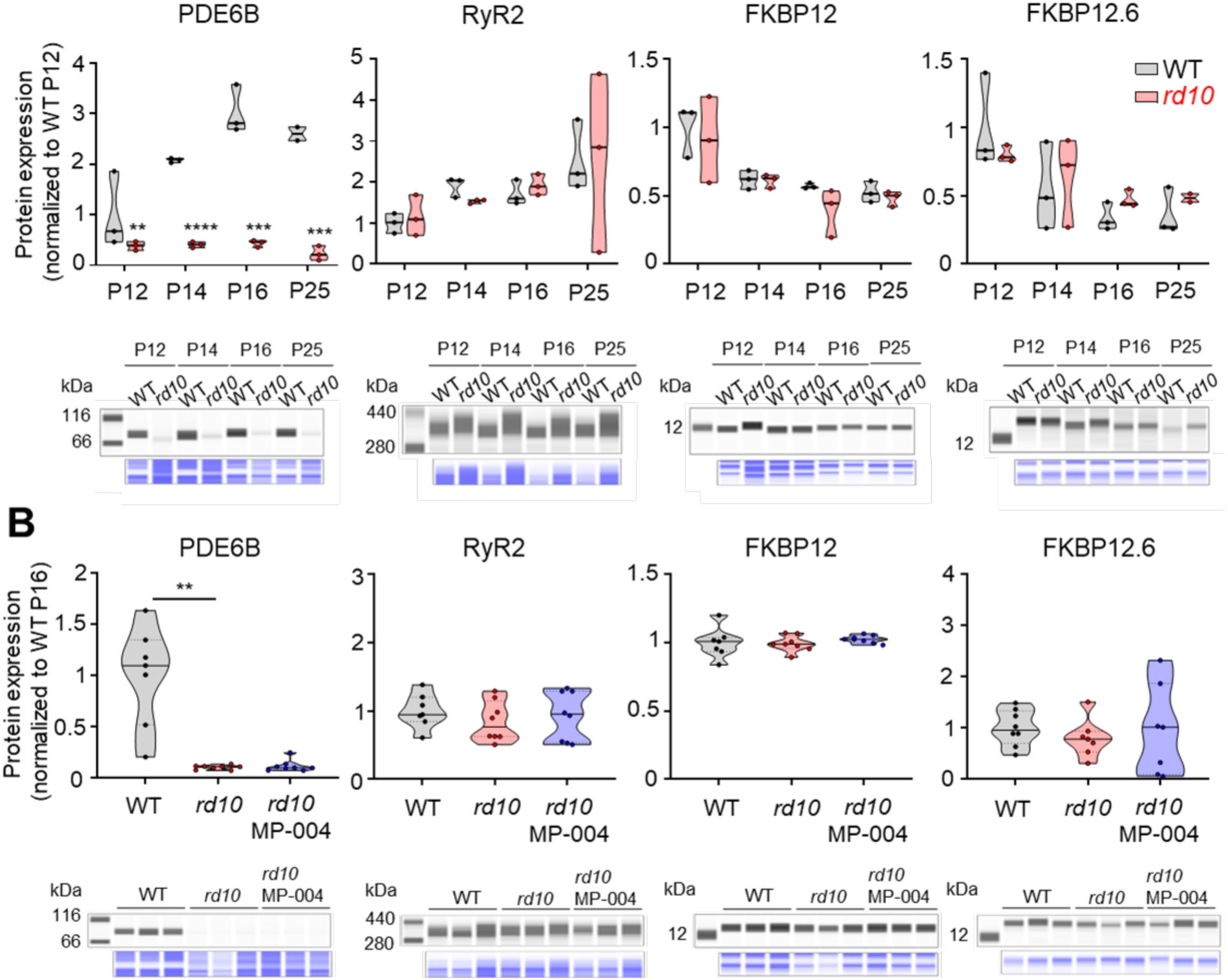
Protein expression in P12-P25 mouse retinas. **A)** Analysis of PDE6B, RyR2, FKBP12 and FKBP12.6 protein levels in WT and *rd10* mouse retinas. Data are normalized to mean P12 WT values. Dots represent data from individual retinas. **p<0.01, ***p <0.001; ****p<0.0001, unpaired t-test. **B)** Effect of MP-004 on protein expression of target proteins was evaluated in P16 *rd10* mouse. Mice were treated daily with vehicle or MP-004 (15 µg/eye) from P12-P15 via topical administration. Data are normalized to mean WT values and presented as violin plots showing the median and interquartile range. Dots represent values from individual mice. **p <0.01, Kruskal Wallis with Dunn’s post-hoc test.

**Supplementary Figure S4.**
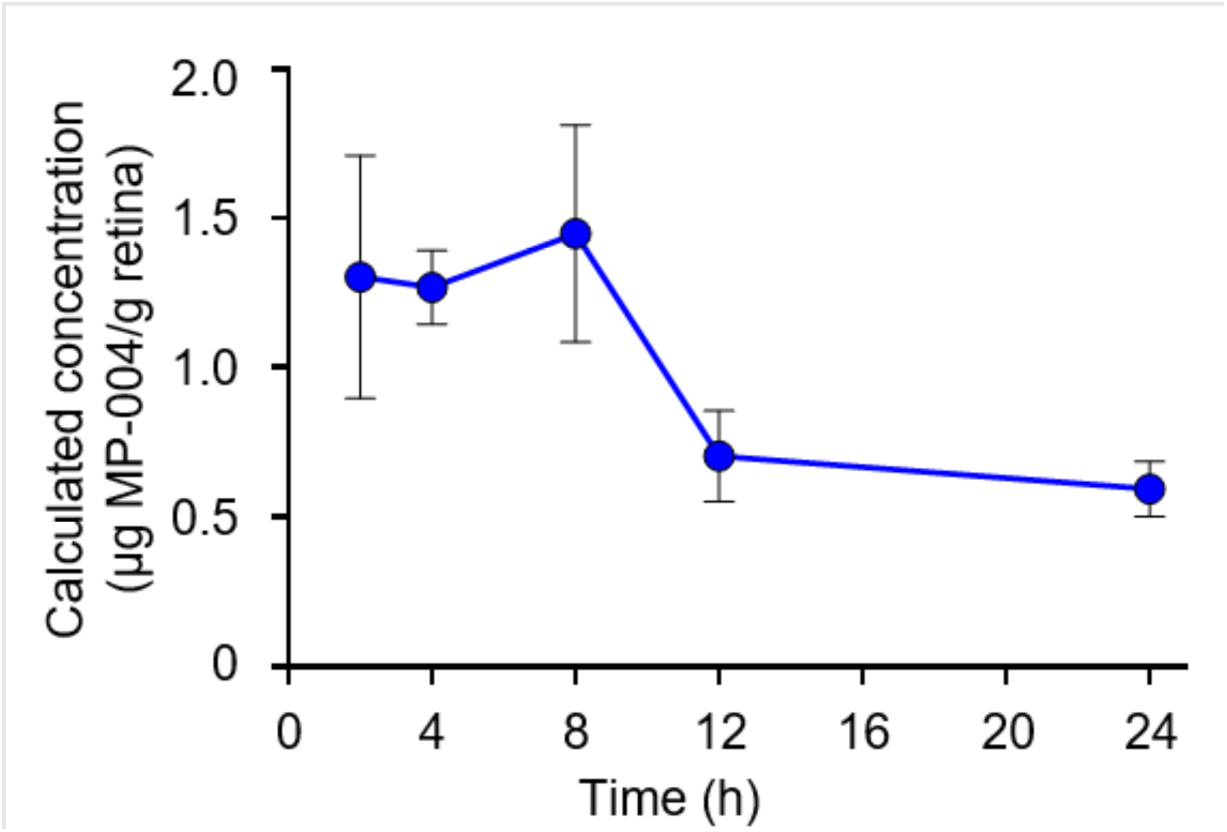
Bioavailability of MP-004 in the mouse retina after ocular instillation (18 µg/eye). Quantitative determination of MP-004 in retina samples was performed by high performance liquid chromatography combined with mass spectrometry (HPLC-MS). MP-004 was detected in retina 24 hours after the topical administration. Data are expressed as mean ± SEM. n=3-6 mice per time point.

**Supplementary Figure S5.**
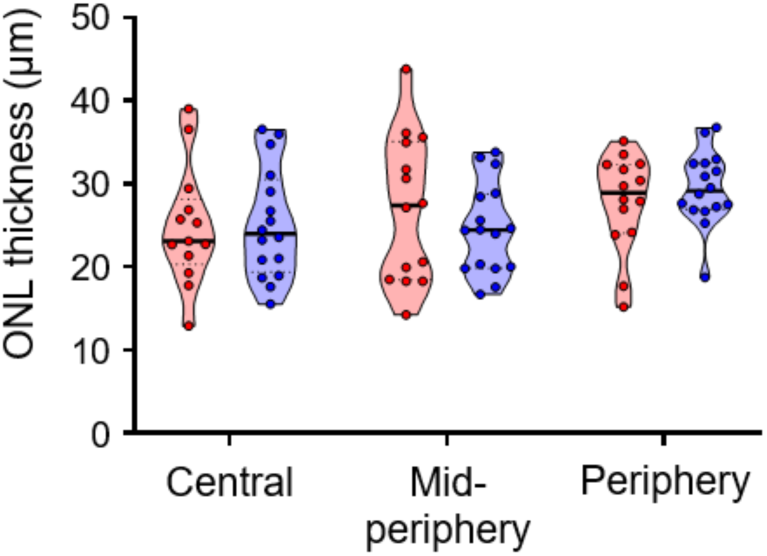
Outer nuclear layer (ONL) thickness of *rd10* retinas treated with vehicle (red) or MP-004 (blue), with at a daily dose of 15 µg per eye. Data are presented as violin plots showing the median and interquartile range. Dots represent values from individual mice.

